# Differential response of patient-derived primary glioblastoma cells to metabolic and adhesion inhibitors

**DOI:** 10.1101/2022.12.19.520994

**Authors:** Rasha Rezk, Fikret Basar, John Mediavillo, Rebecca Donaldson, Colin Watts, Kristian Franze, Alexandre J Kabla

**Affiliations:** York Biomedical Research Institute, Department of Biology, University of York, York, UK; Department of Engineering, University of Cambridge, Cambridge, UK; Birmingham Brain Cancer Program, Institute of Cancer and Genomic Sciences, University of Birmingham, UK; Department of Physiology, Development and Neuroscience, University of Cambridge, Cambridge, UK

## Abstract

**Purpose:** This study aims to investigate Glioblastoma (GBM) cellular response to adhesion and metabolic inhibitors in the context of cells’ migration and cell-matrix adhesion properties. GBM is the most common incurable brain tumour. Decades of work into GBM chemical and molecular classification have identified mechanisms of drug resistance. Inhibitors targeting cancer cell migration and proliferation rarely take into consideration the heterogeneous migration property amongst cells, which may impact patients’ response to treatment.

**Methods:** Tissue samples were obtained from spatially distinct locations with different 5-aminolevulinic acid (5-ALA) fluorescent intensities, strong strongly fluorescent tumour cores, a weak fluorescent tumour rim, and nonfluorescent tumour margins. Samples were previously shown to be associated with different motility and adhesion properties. We tested tumour cells’ response to adhesion and metabolic inhibitors using metabolic assays. Cell survival was also monitored using time-lapse microscopy, while cultured on low-modulus polydimethylsiloxane representative of the stiffness of brain tissue.

**Results:** Metabolic viability assays, MTT and Cell Titer, showed substantial heterogeneity in drug potency. Highly fluorescent tumour core cells were significantly more resistant to an F0F1 ATP synthase inhibitor (Gboxin), and a FAK inhibitor (GSK2256098), and cell proliferation ceased post-treatment in vitro. Cells derived from non-fluorescent tumour margins exhibited higher potency for the ATP synthase inhibitor (Gboxin). However, cell proliferation persisted post-treatment.

**Conclusion:** Our study suggests that the adhesive and migration properties of cells account for the sensitivity to therapeutics in different regions of the tumour in individual patients and between patients with GBM.

## Introduction

Glioblastoma (GBM) is comprised of genetically [1–3] and physically [4] heterogeneous cell populations, which exists within and across patient tumours, presenting a considerable challenge for therapeutic intervention. Genomic and molecular classification of GBM patients have identified four main genomic subtypes, proneural, neural, classical, and mesenchymal [5, 6]. Despite this, it remains unclear whether the different genomic subtypes can predict patient response to anti-angiogenic treatments, such as kinase and proteasome inhibitors. Moreover, distinguishing GBM molecular subtypes has not yet let to improved patient outcomes [7].

Differential response to treatment has also been explored in the context of specific genetic mutations and structural aberrations. A recent pharmacological and genomic profiling of one hundred 100 patient derived GBM cell cultures exposed to 1,544 drugs showed that the differential response to proteasome inhibitors could be linked to mutually exclusive aberrations in TP53 and CDKNK2A/B [8]. Mutant isocitrate dehydrogenase (mtIDH1) also appears to be predictive of treatment response. However, the roles of mutant IDH in cancer development and progression are possibly temporary or dynamic, making the timing of treatment with IDH inhibitors of crucial importance[9].

Preclinical studies mainly focus on the inter-turmour heterogeneity across GBM patients, or a specific gene mutation which are mainly quantified in samples obtained from the resected tumour mass. However, GBM intratumoural molecular heterogeneity strongly limits the performance of therapies targeting specific mutations or molecular subtypes [7]. Conversely, the failure of current therapies to eliminate GBM subpopulations surrounding the edge of the tumour is considered the major factor contributing to the inevitable recurrence. The spatial heterogeneity within GBM tumours have been recently found to facilitate therapeutic resistance [10], where edge-derived cells show a higher capacity for infiltrative growth, while core cells demonstrate core lesions with greater therapy resistance. Understanding GBM intratumour heterogeneity is the key to understanding treatment failure [11]

Intratumour heterogeneity for treatment stratification can be visualised during fluorescent-guided surgery. When 5-aminolevulinic acid (5-ALA) is administered orally prior to surgery, GBM cells glow fluorescent pink, which facilitates maximal resection [12]. However, the level of fluorescence emission varies across the tumour, which may be due to histological and genetic profiling of the tumour [13]. The heterogeneity of 5-ALA-induced fluorescence observed during surgery was associated with different cellular functions and a distinct mRNA expression profile; nonfluorescent tumour tissue resembled the neural subtype of GBM, and fluorescent tumour tissue did not exhibit a pattern that correlates with a known subtype [2]. Arguably, the reduction of 5-ALA fluorescence is suggestive of healthy tissue [14]. Regional variation of biomechanical properties within a tumour could influence therapeutic response and may be a factor that influence the fluorescence of 5-ALA acid-treated cells.

Recently, we demonstrated that tissue fluorescence and spatial heterogeneity is mirrored by physical heterogeneity among GBM primary cells [15]. Cells derived from weak and nonfluorescent tumour rim were smaller, adhered less well, and migrated quicker than cells derived from strongly fluorescent tumour mass. However, whether these properties can predict cellular response treatments remains unclear. To investigate this hypothesis, we tested tumour cells response to two promising adhesion and metabolic inhibitors.

Primary cells were treated with GSK2256098/GTPL7939 (GSK) an ATP competitive reversible inhibitor of focal adhesion kinase (FAK) [16], and Gboxin, a metabolic inhibitor that selectively and irreversibly affects oxygen consumption in GBM cells [17]. GSK is currently in phase II clinical trials for solid cancers treatments such as adenocarcinoma, intracranial and recurrent meningioma mesothelioma and pancreatic cancer [18–23]. FAK is a non-receptor tyrosine kinase which is highly expressed in the nervous system [24] and is activated by cell-extracellular matrix adhesion [25]. FAK is known to play a crucial role in solid cancer progression [26–28]. FAK inhibitors that can selectively bind to the FAK ATP-binding domain are considered the most promising molecules to be translated and applied in clinical practice [29]. The ATP-competitive molecules bind to the FAK–kinase domain competing with ATP and, therefore, inhibiting FAK signal transduction activity and the activation of several FAK downstream pathways [30].

Gboxin is an inhibitor of F0F1 ATP synthase, which plays an important role in cancer metabolism including GBM [31]. A variety of promising F0F1 ATP synthase inhibitors have been reported [32–35], including Bedaquiline which is currently FDA approved [36]. Such inhibitors can offer targeted therapies, by targeting the oxidative phosphorylation pathway in the mitochondria which are dysfunctional in cancer cells [31]. Compared to normal cells, GBM cell have increased mitochondrial membrane potential. Gboxin spares normal cells and selectively targets GBM cells because of the loss of function of mitochondrial permeability transition pore in the cancer cells [17].

We demonstrate a substantial differential response to both treatments across different samples. Cells derived from highly fluorescent tumour core are significantly more resistant to Gboxin, and GSK, compared to cells derived from weak fluorescent tumour rim and nonfluorescent margins that were less adherent with highly migratory behaviour. Highly fluorescent cell proliferation ceased post treatment in vitro when the ATP synthase inhibitor, Gboxin, was administered. While cells derived from non-florescent tumour margins exhibited higher potency for Gboxin, cell proliferation persisted post treatment. Our study suggests that the adhesive and migration properties of cells may contribute to variable sensitivity to both therapeutics in different regions of the tumour in individual patients and between patients with GBM. Such heterogeneity may explain recently observed differential responses of patients to adhesion-blocking drugs.

## Materials and Method

### Sample Collection

Tissue collection protocols complied with the UK Human Tissue Act 2004 (HTA license ref. 12315) and have been approved by the local regional ethics committee (LREC ref. 04/Q0108/60). Tissue samples were derived as described in [15]. Briefly, tissues were derived from newly diagnosed GBM patients who underwent their first surgical resection at Addenbrooke’s, Cambridge University Hospitals. 5-ALA fluorescence was orally administered 4 h before induction of anaesthesia at a dosage of 20 mg/kg. Three different regions within the tumour were biopsied. Six tissue samples from 2 different patients were taken from spatially separated sections using MRI stealth imaging. Navigated biopsy samples were collected from strongly fluorescent tumour cores, a weak fluorescent tumour rim, and nonfluorescent tumour margins. The patient’s clinical information and their molecular biomarker status (IDH mutation and MGMT promoter methylation) can be found in Supplementary Table S1.

### Derivation of GBM Stem-Like Cells

Cell derivation and maintenance follows the protocols described in [37]. The tissue was mechanically minced and enzymatically dissociated before passing through a 40 µm cell strainer. Cells were seeded in serum-free medium (SFM; phenol red-free Neurobasal A) with 2 mM l-glutamine and 1% volume/volume (v/v) penicillin/streptomycin (PS) solution with 20 ng/mL human epidermal growth factor, 20 ng/mL zebrafish fibroblast growth factor (FGF-2), 1% v/v B27 SF supplement, and 1% N2 SF supplement. Cells were allowed to form primary aggregates. Spheroid aggregates were collected and plated onto Engelbreth-Holm-Swarm sarcoma extracellular matrix (ECM, Sigma)–coated flasks (ECM 1:50 dilution with HBSS) and allowed to form a primary monolayer. When the primary monolayer reached 80% confluency, cells were passaged to generate the subsequent monolayers by mechanically and enzymatically dissociating remaining aggregates. Cells were maintained at 37°C and 5% CO2. Experiments were performed using passages 3–9, unless otherwise states. Cell lines were screened regularly for mycoplasma.

### Cytotoxicity Assays

Six thousand cells were cultured in 96 well plates (60 inner wells). Cells were seeded in Triplicates. Following overnight incubation, cells were treated with increasing concentration of Gboxin (Cyman chemical) and GSK2256098 (Cayman Chemical) as indicated in the corresponding figure legends.

Gboxin viability assay was performed 40-42 hours after treatment as per the protocol provided by manufacturer for Cell Titer Glo® (Promega). GSK viability assay was performed 18 hours after treatment as per the protocol provided by the manufacturer for Thiazolyl blue tetrazolium bromide (Sigma).

Blanks (outer wells, without cells) were included during the cell titer and MTT incubation for proper background subtraction. Data are expressed as a percentage of viability when compared with untreated cells (negative control—considered as 100% of viability). For the normalisation of the data, treated cells were subtracted from cells treated with the corresponding dimethyl sulfoxide (DMSO) concentration present in each drug concentration. The 50% inhibitory concentration (IC_50_) for each drug was defined as the concentration producing 50% less viability comparted to control wells.

### Polydimethylsiloxane (PDMS) Substrates for Studying Cellular Motility and Morphology Post-treatment

Substrates were prepared as described in [15]. NuSil GEL-8100 (NuSil) was prepared in a 1:1 ratio of component A and component B and mixed well for 60 seconds: 1% (w/w) 10:1 (base/crosslinker w/w) Sylgard-184 (VWR) was added to the GEL-8100 and mixed well for 60 seconds. 80 mg per well of PDMS was added to 24-well culture plates (Corning Life Science). Coated vessels were baked at 65°C for 13 h. This treatment gave a shear modulus value of G = 1.53 ± 0.12 kPa. Ten thousand cells were cultured in 24 well plates. Following overnight incubation, cells were treated with Gboxin and GSK concentration which correspond to 50% ± 3% viability for the given cell line. Cells were seeded in triplicates and imaged every 10 min for 48 hr. Time-lapse images were acquired with Zeiss Axio Observer Z1.

### Statistical Analysis

Data were collected from at least 3 independent biological experiments. The order of data collection was randomized; no blinding was performed and no data were excluded from the analysis.

Representative results for biological replicates are shown for every figure except where specified otherwise in the figure legends. Data are presented as mean ± SEM as specified in the figure legend. Cell viability analysis was performed using software GraphPad Prism 7.0a.

Statistical significance with exact p value was determined with the method used as indicated in corresponding figure legend. First, we tested differences between patients and differences between fluorescence groups independently. Then we tested for differences simultaneously using an additive model (for patients and fluorescence intensity). There was no statistically significant difference between weak- and nonfluorescent cell lines but both were different from strongly fluorescent cell lines. To check whether sampling from multiple locations violated our standard linear model assumptions, we tested unexplained heterogeneity using mixed-effect regression. Including a random effect of all lines did not significantly improve model fit. This indicates that a mixed effect model is not required. Statistical analyses and plotting were performed using the R statistical software package.

## Results

### Gboxin and GSK toxicity in GBM cells

To investigate GBM cellular response to adhesion and metabolic inhibitors, we tested different cell populations derived from different GBM tumours, as well as different regions from within the tumours (Figure 1A). To understand the variability in cell toxicity observed between patient cell lines (Figure 1B, Figure 1C), we first tested drug potency (IC50 values) across different patients. Cells derived from patient A were less sensitive to GSK (***P < .0001) and Gboxin (*P < 0.01) compared to cells derived from patient B. There was no statistically significant difference between weak- and nonfluorescent cell lines response to Gboxin, but both were different from strongly fluorescent cell lines (*P < 0.01). We adjusted our model to account for fluorescence intensity when comparing between patients (Methods). We found that cells from weak- and non-fluorescent tumour cells derived from patient A and patient B had significantly higher toxicity to both drugs compared to cells strongly fluorescent core samples (***P < .0001).

**Figure 1:**
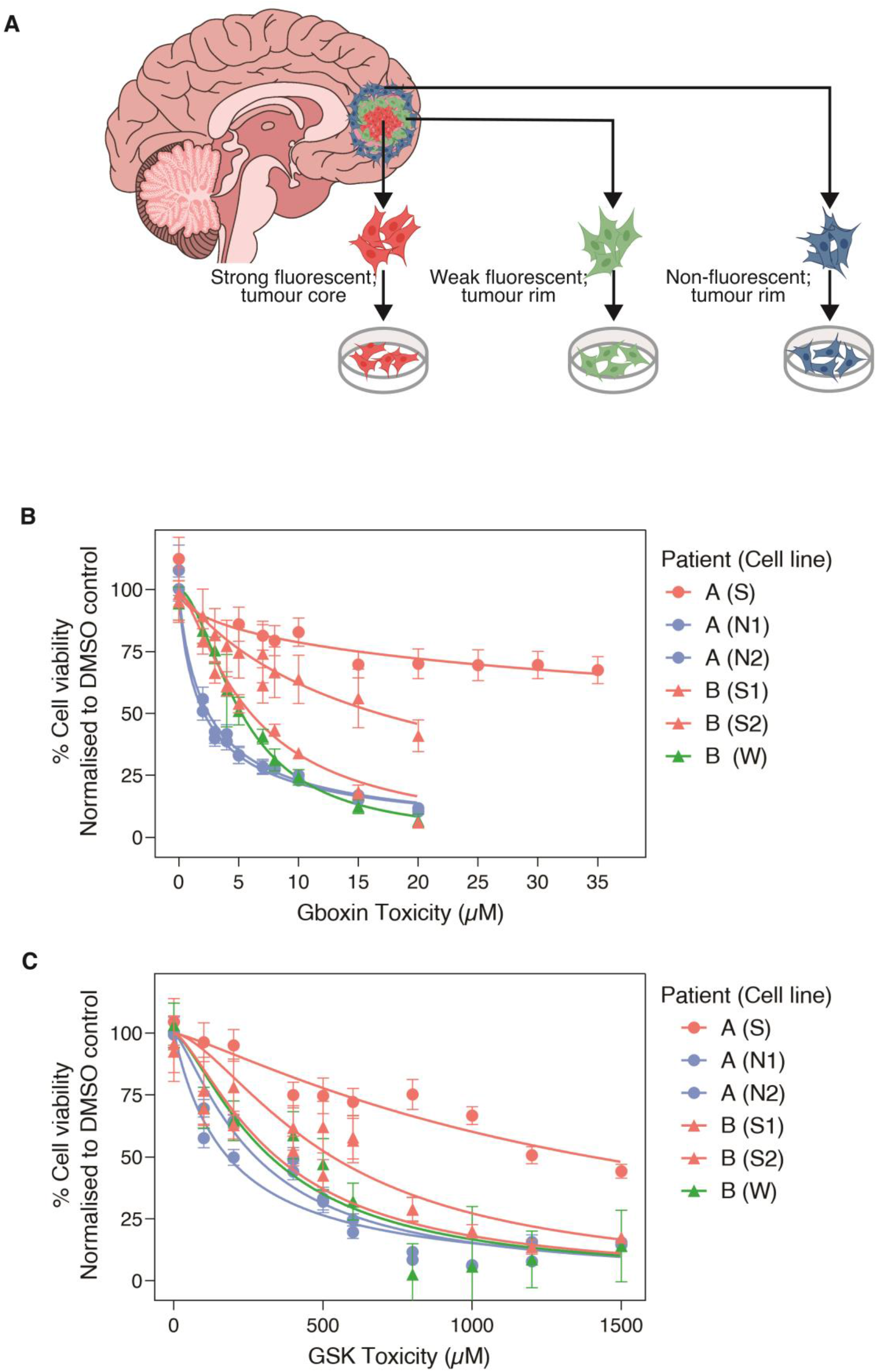
GBM cells exhibit differential treatment response to metabolic and adhesion inhibitors. A) We collected three strongly fluorescent tissue samples derived from the tumour mass (S from patient A, S1 and S2 from patient B, data shown in red on graphs) two nonfluorescent tumour margins (N1 and N2 from patient A, data in blue), and 1 weak fluorescent tissue sample (W from patient B, data in green). B) Cell viability as a function of Gboxin concentration. Error bars represent mean ± SEM from three independent experiments. C) Cell viability as a function of GSK concentration. Error bars represent mean ± SEM from three independent experiments.

The results demonstrate viability differences within each tumour and between tumours and show that this heterogeneity is related to and 5-ALA fluorescence intensity.

### Gboxin does not disrupt cell proliferation for cells derived from nonfluorescent tumour margins

To determine whether GSK and Gboxin will achieve the desired primary effect *in vitro*, we observed cell viability and proliferation on a compliant PDMS substrate, before treatment, for 24 hours, and after administering GSK and Gboxin (IC50 ± 3 %) for 48 hours. Prior to treatments cell division/proliferation was observed across all cell lines (n=3) (Supplementary video 1, video 2 and video 3). Around 65% of GSK treated cells lost their shape, detached, and become afloat, within 18 hours (n=3), reaching 90% within 48 hours (n=3) (Figure 2). No cell division or proliferation was observed across non-fluorescent, weak and strong fluorescent cell lines for 48 hours post GSK treatment (Supplementary video 4, video 5, video 6).

**Figure 2:**
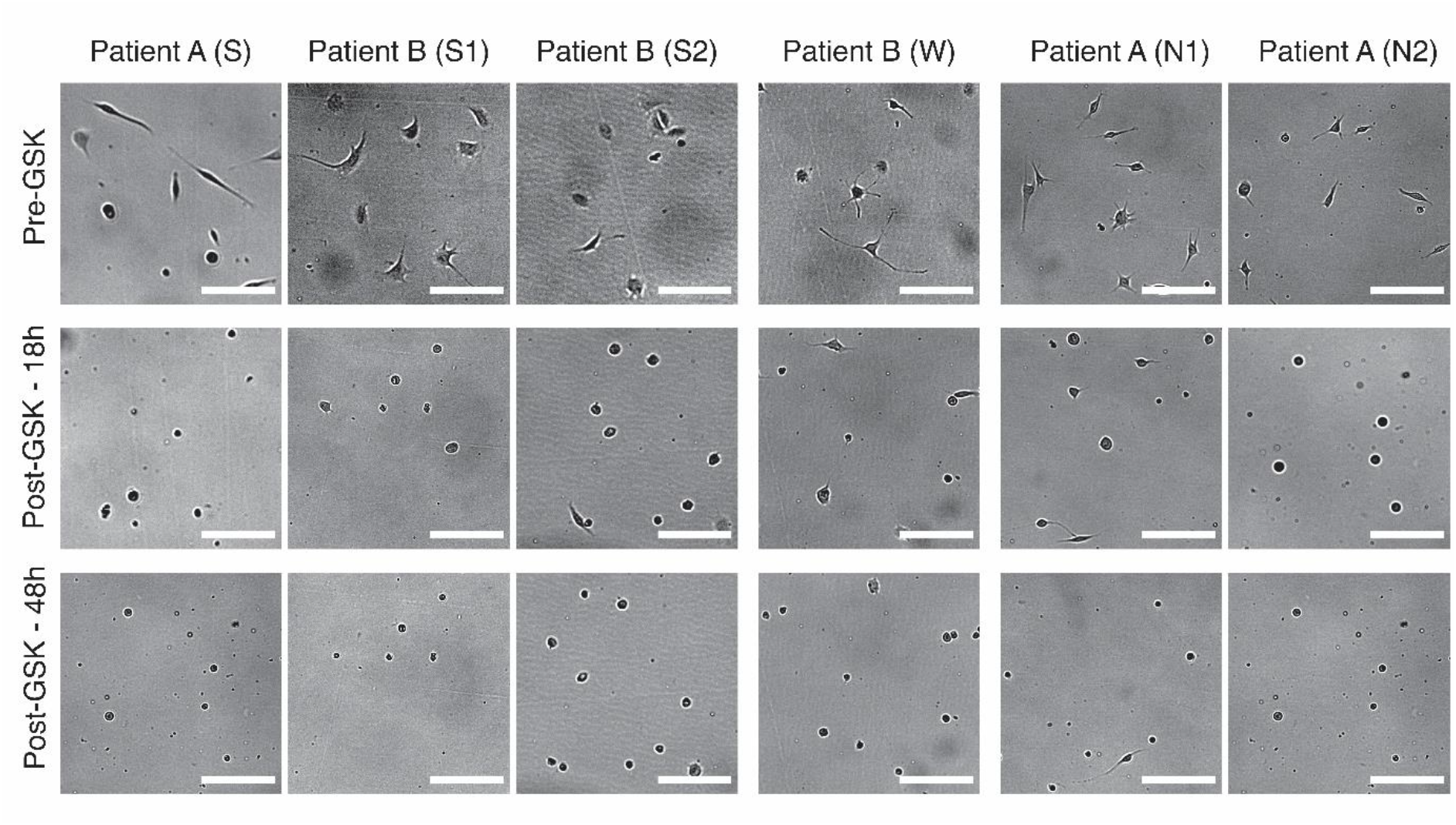
Cytotoxicity effect of GSK

**Figure 3:**
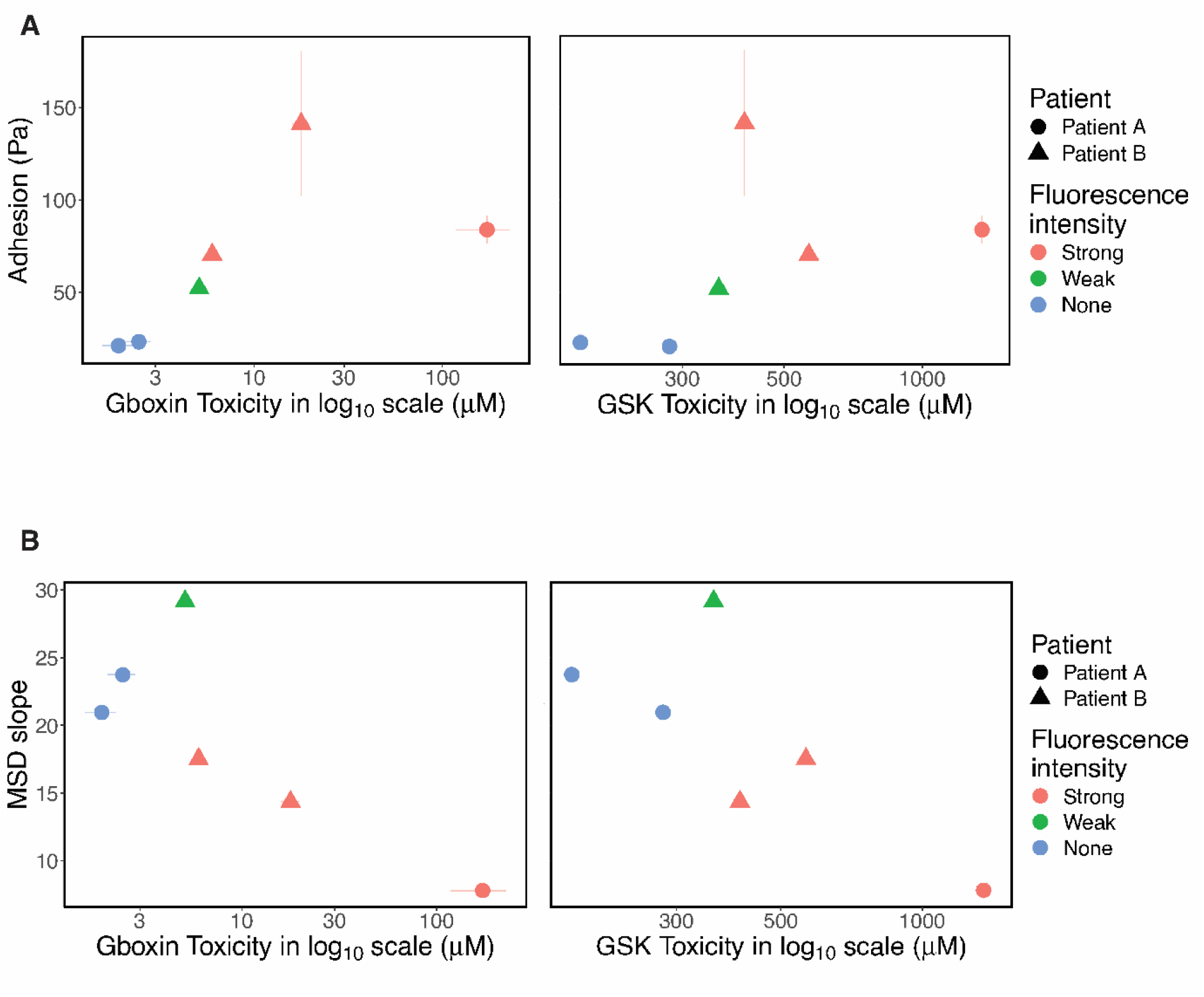
Cell-matrix adhesion and cell migration are important factors for GBM treatment response. A) Relationship between drug potency (IC50) and cell-matrix adhesion strength. Strong fluorescent cells are more adherent and are more resistant to both Gboxin and GSK. Error bars represent SEM from at least 2 independent experiments. B) Relationship between drug potency (IC50) and cell migration. Strong fluorescent cells are more adherent and are more resistant to both Gboxin and GSK. Error bars represent SEM from at least 2 independent experiments.

Around 62% of Gboxin cells died/lost their shape, detached, and become afloat within 42 hours. No cell division was observed across cells derived from strong fluorescent tumour core and weak florescent tumour rim (for both patients, Supplementary video 7, video 8). However, cells derived from nonfluorescent tumour margins (both cell lines) proliferated/divided (Supplementary video 9, video 10).

The N2 cell line seemed to have a higher rate of proliferation compared to the N1 cell line post Gboxin treatment, this may have been due to the higher relative Gboxin concentration administered to N1 (corresponding to 42.7% and 49.2% cell viability for N1 and N2 respectively).

All primary cells were treated for 48 h with the corresponding IC50. Over 90% cells lost their shape, become round and, detached become afloat.

The results establish/reinforce that cells derived from highly fluorescent tumour core are significantly more resistant (an order of magnitude) to GSK and Gboxin treatments, compared to cells derived from weak fluorescent tumour rim and nonfluorescent tumour margins that were less adherent with highly migratory behaviour. However, cells derived from nonfluorescent tumour margins retain their proliferation after administrating Gboxin in culture.

## Discussion

Differential response of GBM to anti-angiogenic treatments, such as kinase and proteasome inhibitors, represents a challenge for therapeutic development. Factors associated with patients’ differential response remains unclear. While genomic and transcriptomic profiling help categorise GBM tumours into different subtypes [5], they have not led to improvement of patients’ outcomes due to the complexities arising from intratumoural molecular heterogeneity [7]. Intratumour heterogeneity remains a major obstacle to therapeutic intervention.

Our results reveal that Gboxin potency inversely correlates with cell migration which is also associated with 5-ALA intensity. The spatial distinction is also in agreement with a recent study, by Bastola et al., which showed that intratumour spatial heterogeneity facilitates therapeutic resistance where edge-derived cells show a higher capacity for infiltrative growth, and core cells demonstrated core lesions with greater therapy resistance in xenotransplantation, and invitro radiation therapy [10].

A possible explanation lies in the Warburg effect, in which highly proliferative cells undergo a metabolic shift from oxidative phosphorylation, which is targeted by Gboxin, to aerobic glycolysis [38]. GBM intratumoural cell proliferation is strongly associated with the fluorescence intensity of samples and hence the spatial density and distribution of cells. Glioma cells with higher 5-ALA fluorescence were shown to be more proliferative than those with lower fluorescence [12]. This could explain the Gboxin resistance exhibited by the strongly fluorescent, GBM tumour core cells in our study. In addition, this metabolic alteration is characterised by an increase in the ratio of oxidised and reduced forms of the carrier molecule Nicotinamide adenine dinucleotide (NAD+/NADH) [39, 40]. Yamashita et al showed this ratio is higher at the core of GBM tumours than the periphery [41], implying a shift to aerobic glycolysis in the tumour core. Interestingly however, the assuming less proliferative cell lines (non-fluorescent tumour margins), while they were more sensitive to Gboxin, their proliferation was not disturbed by the drug in culture.

Our earlier work demonstrated links between fluorescence levels and mechanical attributes of the cells. The distinct adhesion and migratory profiles of GBM suggests that patients’ differential response to adhesion inhibitors and anti-invasive molecular treatments may be due to intrinsic differences in cell motility, adhesion, and traction forces. We therefore expect Gboxin and GSK toxicity to show similar correlations with cell-matrix adhesion and migratory properties.

Cells derived from tumour core with the highest cell-adhesion strength exhibited the highest resistance to both treatments, GSK and Gboxin. Cells derived from the weak fluorescent tumour rim and the non-fluorescent tumour margin which all had significantly lower adhesion properties (compared to strong fluorescent core) had much higher potency to both drugs, while cell migration inversely correlated with both drug potency.

As expected, cells treatment response to GSK directly correlated with cell-ECM adhesion properties, which was also associated with the amount of 5-ALA fluorescence, and cell migration. Cell-ECM adhesion plays a role in cellular migration, invasion [42], and proliferation [43]. Highly adhesive, strongly fluorescent GBM stem cells were less potent to GSK. This can be explained as FAK has a fundamental role in adhesion and is recruited to stabilise focal adhesions [44]. Therefore, more adhesive cells are likely to have more FAK and a larger cytotoxic effect when FAK is inhibited. This increased cytotoxic effect is explained by the fact that when FAK is inhibited, the EGFR signalling pathway is inhibited at the cellular membrane and a pro-apoptotic signal is transmitted to the nucleus [42].

Preclinical studies of kinase and proteasome inhibitors typically use lines from the resected tumour core/mass, which may not be representative of the distribution of the adhesion and motility properties of GBM edge like cells. While GSK and Gboxin potency exhibited an identical/similar pattern across cell lines. Our results show that GSK potency (IC50 values) for cells derived from the strongly fluorescent tumour core is an order of magnitude higher than what is reported when treating pancreatic ductal adenocarcinoma cells [45].

Therapeutic modalities could benefit from targeting the edge-located tumour-initiating cells from GBM patients [10]. Our study reveal that cells derived from both GBM tumour rim and tumour margins have a distinct response to treatment. It also showed that while drug potency was higher for cells derived from the edge located tumour cells, they were still able to proliferate and grow in culture after Gboxin treatment. Our study reveals sampling is crucial and the cell adhesive and motility properties mechanics may play a role in drug sensitive/screening.

## Funding

This research was supported by the University of Cambridge (Engineering for the Clinical Practice grants) to R. R. and A.J.K. R.R. was supported by Nabaa El Mahabaa (Egypt) and Gonville & Caius College.

## Data availability

Research data are stored in an institutional repository and will be shared upon request to the corresponding author.

## Author contribution

R.R. conceptualized and designed the study. R.R, F.B., J.M. and R.D. and A.J.K designed and performed the experiments, and analysed the data. C.W. supervised the clinical procedure, provided surgical samples, and interpreted clinical data. R.R, F.B., J.M., K.F. and A.J.K discussed the data and wrote the manuscript.

## Conflict of interest

The authors report no conflict of interest, and the present study has not been presented elsewhere.

---

**Supplementary Table 1:** Patients Information and histological analysis

| Patient | Age | Sex | MGMT | IDH1 | MIB-1 (Ki-67%) |
| --- | --- | --- | --- | --- | --- |
| Patient A | 61 | Male | Unmethylated | wt | 21 |
| Patient B | 67 | Male | Methylated | wt | 13.2 |
| Patient C | 54 | Male | Unmethylated | wt | 40 |

## Supplementary Video

1. Cells viability and proliferation observed on a compliant PDMS substrate prior to Gboxin treatment. Cells were derived from non-fluorescent tumour rim
2. Cells viability and proliferation observed on a compliant PDMS substrate prior to GSK treatment. Cells were derived from weak tumour trim.
3. Cells viability and proliferation observed on a compliant PDMS substrate, before prior to GSK treatment. Cells were derived from strong tumour core
4. Cells viability and proliferation observed on a compliant PDMS substrate post to GSK treatment. Cells were derived from non-fluorescent tumour margin
5. Cells viability and proliferation observed on a compliant PDMS substrate post to GSK treatment. Cells were derived from weak tumour trim.
6. Cells viability and proliferation observed on a compliant PDMS substrate, before post to GSK treatment. Cells were derived from strong tumour core
7. Cells viability and proliferation observed on a compliant PDMS substrate, before post to Gboxin treatment. Cells were derived from weak tumour rim
8. Cells viability and proliferation observed on a compliant PDMS substrate, before post to Gboxin treatment. Cells were derived from strong tumour core
9. Cells viability and proliferation observed on a compliant PDMS substrate, before post to Gboxin treatment. Cells were derived from non-fluorescent tumour marhin
10. Cells viability and proliferation observed on a compliant PDMS substrate, before post to Gboxin treatment. Cells were derived from non-fluorescent tumour margin

## Notes

### Competing Interest Statement

The authors have declared no competing interest.

## References

1. Bonavia R, Mukasa A, Narita Y, Sah DW, Vandenberg S, Brennan C, Johns TG, Bachoo R, Hadwiger P, Tan P (2010) Tumor heterogeneity is an active process maintained by a mutant EGFR-induced cytokine circuit in glioblastoma. Genes & development 24: 1731–1745

2. Almiron Bonnin DA, Havrda MC, Lee MC, Evans L, Ran C, Qian DC, Harrington LX, Valdes PA, Cheng C, Amos CI, Harris BT, Paulsen KD, Roberts DW, Israel MA (2019) Characterizing the heterogeneity in 5-aminolevulinic acid-induced fluorescence in glioblastoma. J Neurosurg 132: 1706–1714 doi:10.3171/2019.2.Jns183128

3. Patel AP, Tirosh I, Trombetta JJ, Shalek AK, Gillespie SM, Wakimoto H, Cahill DP, Nahed BV, Curry WT, Martuza RL (2014) Single-cell RNA-seq highlights intratumoral heterogeneity in primary glioblastoma. Science 344: 1396–1401

4. Foss A, Zanoni M, So WY, Jenkins L, Tosatto L, Bartolini D, Gottesman MM, Tesei A, Tanner K (2020) Patient-derived glioblastoma cells (GBM) exhibit distinct biomechanical profiles associated with altered activity in the cytoskeleton regulatory pathway. BioRxiv

5. Verhaak RG, Hoadley KA, Purdom E, Wang V, Qi Y, Wilkerson MD, Miller CR, Ding L, Golub T, Mesirov JP (2010) Integrated genomic analysis identifies clinically relevant subtypes of glioblastoma characterized by abnormalities in PDGFRA, IDH1, EGFR, and NF1. Cancer cell 17: 98–110

6. Brennan CW, Verhaak RG, McKenna A, Campos B, Noushmehr H, Salama SR, Zheng S, Chakravarty D, Sanborn JZ, Berman SH (2013) The somatic genomic landscape of glioblastoma. Cell 155: 462–477

7. Lee E, Yong RL, Paddison P, Zhu J Comparison of glioblastoma (GBM) molecular classification methods. Seminars in cancer biology. Elsevier, pp 201–211

8. Johansson P, Krona C, Kundu S, Doroszko M, Baskaran S, Schmidt L, Vinel C, Almstedt E, Elgendy R, Elfineh L, Gallant C, Lundsten S, Ferrer Gago FJ, Hakkarainen A, Sipilä P, Häggblad M, Martens U, Lundgren B, Frigault MM, Lane DP, Swartling FJ, Uhrbom L, Nestor M, Marino S, Nelander S (2020) A Patient-Derived Cell Atlas Informs Precision Targeting of Glioblastoma. Cell Reports 32: 107897 doi:https://doi.org/10.1016/j.celrep.2020.107897

9. Pirozzi CJ, Yan H (2021) The implications of IDH mutations for cancer development and therapy. Nature Reviews Clinical Oncology 18: 645–661

10. Bastola S, Pavlyukov MS, Yamashita D, Ghosh S, Cho H, Kagaya N, Zhang Z, Minata M, Lee Y, Sadahiro H, Yamaguchi S, Komarova S, Yang E, Markert J, Nabors LB, Bhat K, Lee J, Chen Q, Crossman DK, Shin-Ya K, Nam D-H, Nakano I (2020) Glioma-initiating cells at tumor edge gain signals from tumor core cells to promote their malignancy. Nature Communications 11: 4660 doi:10.1038/s41467-020-18189-y

11. Sottoriva A, Spiteri I, Piccirillo SG, Touloumis A, Collins VP, Marioni JC, Curtis C, Watts C, Tavaré S (2013) Intratumor heterogeneity in human glioblastoma reflects cancer evolutionary dynamics. Proceedings of the National Academy of Sciences 110: 4009–4014

12. Widhalm G, Wolfsberger S, Minchev G, Woehrer A, Krssak M, Czech T, Prayer D, Asenbaum S, Hainfellner JA, Knosp E (2010) 5-Aminolevulinic acid is a promising marker for detection of anaplastic foci in diffusely infiltrating gliomas with nonsignificant contrast enhancement. Cancer: Interdisciplinary International Journal of the American Cancer Society 116: 1545–1552

13. Mazurek M, Szczepanek D, Orzyłowska A, Rola R (2022) Analysis of Factors Affecting 5-ALA Fluorescence Intensity in Visualizing Glial Tumor Cells-Literature Review. Int J Mol Sci 23: 926 doi:10.3390/ijms23020926

14. Della Puppa A, Munari M, Gardiman MP, Volpin F (2019) Combined fluorescence using 5-aminolevulinic acid and fluorescein sodium at glioblastoma border: intraoperative findings and histopathologic data about 3 newly diagnosed consecutive cases. World neurosurgery 122: e856–e863

15. Rezk R, Jia BZ, Wendler A, Dimov I, Watts C, Markaki AE, Franze K, Kabla AJ (2020) Spatial heterogeneity of cell-matrix adhesive forces predicts human glioblastoma migration. Neurooncol Adv 2: vdaa081–vdaa081 doi:10.1093/noajnl/vdaa081

16. Brown NF, Williams M, Arkenau H-T, Fleming RA, Tolson J, Yan L, Zhang J, Singh R, Auger KR, Lenox L (2018) A study of the focal adhesion kinase inhibitor GSK2256098 in patients with recurrent glioblastoma with evaluation of tumor penetration of [11C] GSK2256098. Neuro-oncology 20: 1634–1642

17. Shi Y, Lim SK, Liang Q, Iyer SV, Wang H-Y, Wang Z, Xie X, Sun D, Chen Y-J, Tabar V, Gutin P, Williams N, De Brabander JK, Parada LF (2019) Gboxin is an oxidative phosphorylation inhibitor that targets glioblastoma. Nature 567: 341–346 doi:10.1038/s41586-019-0993-x

18. ClinicalTrials.gov [Internet] (2015) Positron Emission Tomography (PET) Study to Evaluate Biodistribution of 11C-GSK2256098 in Healthy Subjects and Idiopathic Pulmonary Arterial Hypertension (PAH) Patients. National Library of Medicine (US). 2000 Feb 29. https://ClinicalTrials.gov/show/NCT02551653. Accessed November 21 2022

19. https://ClinicalTrials.gov [Internet] (2015) A Study of GSK2256098 and Trametinib in Advanced Pancreatic Cancer. National Library of Medicine (US). 2000 Feb 29. https://ClinicalTrials.gov/show/NCT02428270. Accessed November 21 2022

20. ClinicalTrials.gov [Internet] (2013) A Dose Escalation Study to Assess Safety of GSK2256098 (FAK Inhibitor) in Combination With Trametinib (MEK Inhibitor) in Subjects With Advanced Solid Tumors. National Library of Medicine (US). 2000 Feb 29. https://ClinicalTrials.gov/show/NCT01938443. Accessed November 21 2022

21. ClinicalTrials.gov [Internet] (2010) Study of a Focal Adhesion Kinase Inhibitor in Subjects With Solid Tumors. National Library of Medicine (US). 2000 Feb 29. https://ClinicalTrials.gov/show/NCT01138033. Accessed November 21 2022

22. ClinicalTrials.gov [Internet] (2009) Phase I Study to Evaluate Safety, Pharmacokinetics and Pharmacodynamics of GSK2256098 in Healthy Volunteers. National Library of Medicine (US). 2000 Feb 29. https://ClinicalTrials.gov/show/NCT00996671. Accessed November 21 2022

23. ClinicalTrials.gov [Internet] (2015) Vismodegib, FAK Inhibitor GSK2256098, Capivasertib, and Abemaciclib in Treating Patients With Progressive Meningiomas. National Library of Medicine (US). 2000 Feb 29. https://ClinicalTrials.gov/show/NCT02523014. Accessed November 21 2022

24. Contestabile A, Bonanomi D, Burgaya F, Girault J-A, Valtorta F (2003) Localization of focal adhesion kinase isoforms in cells of the central nervous system. International Journal of Developmental Neuroscience 21: 83–93 doi:https://doi.org/10.1016/S0736-5748(02)00126-0

25. Xiong W-C, Mei L (2003) Roles of FAK family kinases in nervous system. Frontiers in Bioscience-Landmark 8: 676–682

26. Perrone C, Pomella S, Cassandri M, Braghini MR, Pezzella M, Locatelli F, Rota R (2020) FAK Signaling in Rhabdomyosarcoma. Int J Mol Sci 21: 8422

27. Aboubakar Nana F, Hoton D, Ambroise J, Lecocq M, Vanderputten M, Sibille Y, Vanaudenaerde B, Pilette C, Bouzin C, Ocak S (2019) Increased Expression and Activation of FAK in Small-Cell Lung Cancer Compared to Non-Small-Cell Lung Cancer. Cancers 11: 1526

28. Tapial Martínez P, López Navajas P, Lietha D (2020) FAK Structure and Regulation by Membrane Interactions and Force in Focal Adhesions. Biomolecules 10: 179

29. Pomella S, Cassandri M, Braghini MR, Marampon F, Alisi A, Rota R (2022) New Insights on the Nuclear Functions and Targeting of FAK in Cancer. Int J Mol Sci 23: 1998

30. Golubovskaya VM (2014) Targeting FAK in human cancer: from finding to first clinical trials. FBL 19: 687–706 doi:10.2741/4236

31. Wu Z, Ho WS, Lu R (2021) Targeting mitochondrial oxidative phosphorylation in glioblastoma therapy. Neuromolecular Medicine: 1–5

32. Fewtrell C, Gomperts B (1977) Quercetin: A novel inhibitors of Ca2+ influx and exocytosis in rat peritoneal mast cells. Biochimica et Biophysica Acta (BBA)-Biomembranes 469: 52–60

33. Datiles MJ, Johnson EA, McCarty RE (2008) Inhibition of the ATPase activity of the catalytic portion of ATP synthases by cationic amphiphiles. Biochimica et Biophysica Acta (BBA)-Bioenergetics 1777: 362–368

34. Blatt NB, Bednarski JJ, Warner RE, Leonetti F, Johnson KM, Boitano A, Yung R, Richardson BC, Johnson KJ, Ellman JA (2002) Benzodiazepine-induced superoxide signalsB cell apoptosis: mechanistic insight and potential therapeutic utility. The Journal of clinical investigation 110: 1123–1132

35. Johnson KM, Chen X, Boitano A, Swenson L, Opipari Jr AW, Glick GD (2005) Identification and validation of the mitochondrial F1F0-ATPase as the molecular target of the immunomodulatory benzodiazepine Bz-423. Chemistry & biology 12: 485–496

36. Fiorillo M, Lamb R, Tanowitz HBB, Cappello ARR, Martinez-Outschoorn UEE, Sotgia F, Lisanti MPP (2016) Bedaquiline, an FDA-approved antibiotic, inhibits mitochondrial function and potently blocks the proliferative expansion of stem-like cancer cells (CSCs). Aging 8: 1593–1606 doi:10.18632/aging.100983

37. Fael Al-Mayhani TM, Ball SL, Zhao JW, Fawcett J, Ichimura K, Collins PV, Watts C (2009) An efficient method for derivation and propagation of glioblastoma cell lines that conserves the molecular profile of their original tumours. J Neurosci Methods 176: 192–199 doi:10.1016/j.jneumeth.2008.07.022

38. Warburg O (1956) On respiratory impairment in cancer cells. Science 124: 269–270

39. Deberardinis RJ, Lum JJ, Hatzivassiliou G, Thompson CB (2008) The biology of cancer: metabolic reprogramming fuels cell growth and proliferation. Cell metabolism 7 1: 11–20

40. da Veiga Moreira J, Hamraz M, Abolhassani M, Bigan E, Pérès S, Paulevé L, Levy Nogueira M, Steyaert J-M, Schwartz L (2016) The redox status of cancer cells supports mechanisms behind the Warburg effect. Metabolites 6: 33

41. Yamashita D, Botta D, Cho HJ, Guo X, Ozaki S, Flanary VL, Sirota I, Gao M, Yamaguchi S, Nakano MA, Zhou F, Zhou H, Kondo T, Kunieda T, Crossman DK, Kornblum HI, Gorospe M, Nam D-H, Zamboni N, Skolnick J, Gu Z, Lund FE, Nakano I (2020) Spatial heterogeneity of glioblastoma cells reveals sensitivity to NAD+ depletion at tumor edge. bioRxiv: 2020.2011.2026.399725 doi:10.1101/2020.11.26.399725

42. Jones G, Machado J, Merlo A (2001) Loss of focal adhesion kinase (FAK) inhibits epidermal growth factor receptor-dependent migration and induces aggregation of NH2-terminal FAK in the nuclei of apoptotic glioblastoma cells. Cancer research 61: 4978–4981

43. Schwartz MA, Assoian RK (2001) Integrins and cell proliferation: regulation of cyclin-dependent kinases via cytoplasmic signaling pathways. Journal of cell science 114: 2553–2560

44. Nader GP, Ezratty EJ, Gundersen GG (2016) FAK, talin and PIPKIγ regulate endocytosed integrin activation to polarize focal adhesion assembly. Nature Cell Biology 18: 491–503

45. Zhang J, He D-H, Zajac-Kaye M, Hochwald SN (2014) A small molecule FAK kinase inhibitor, GSK2256098, inhibits growth and survival of pancreatic ductal adenocarcinoma cells. Cell cycle 13: 3143–3149

